# Idea paper: An envelope model of ecological disturbance

**DOI:** 10.1101/2021.12.11.472203

**Authors:** Nicholas R. Friedman

## Abstract

Disturbance is common in natural ecosystems, but increasingly defines them. While there are many descriptions for the dynamics of an ecosystem’s response to disturbance, there are few descriptions for the dynamics of the disturbance itself. I describe a novel application of a model based on the production of amplitude envelopes in acoustics and electronic music synthesis, with varying parameters Attack, Decay, Sustain, and Release (ADSR). I show that varying the parameters of the ADSR model is sufficient to produce and vary the qualitative disturbance regimes described by previous authors, and is capable of producing dynamics not previously considered. I tested the utility of the ADSR model by applying it to a logistic growth model. I found that manipulating the attack and release parameters of the ADSR model changes the population dynamics estimated by these models. This implies that responses to disturbance are determined not only by the resilience and resistance of the ecological system, but also the dynamics of the disturbance itself. My hope is that the ADSR model will prove useful to researchers in either describing disturbances in long-term ecological data, or in producing disturbances for simulations or experiments.

## Research Question

How does variation in the temporal dynamics of ecological disturbance impact the characteristics of ecological response?

## Value

Humans are altering the natural environment in many ways that perturb the planet’s ecosystems (Vitousek, 1994) or disrupt natural disturbance regimes (Turner, 2010). Ecological responses to disturbance vary widely, and researchers can use measurements such as resilience and resistance to characterize and in principle predict these responses (Donohue et al., 2013). Disturbance also varies widely, and its temporal dynamics can impact the trajectory of ecological responses (Jacquet & Altermatt, 2020). A flexible model for generating or describing disturbance dynamics would contribute to our ability to test and predict the effects of various disturbances on ecological systems.

## Relevant Hypothesis

In his seminal paper, Lake (2000; see also Bender et al., 1984) explored the temporal dynamics of disturbance in floods and droughts, and described three dynamics common in nature: “Pulse” events like floods, which are discrete and short-term, “Press” events like sedimentation, which are maintained at a constant level, and “Ramp” events like droughts, which steadily become more severe. While these categories have helped to shape ecologists’ questions about disturbance (Ryo et al., 2019), many disturbances do not fit these shapes. Indeed, the temporal dynamics of flow rate during floods (Federer et al., 1990), radioactive iodine levels following the great Tohoku earthquake and tsunami (Inui et al., 2012), and atmospheric aerosols following volcanic eruption (Sato et al., 1993) all show a different disturbance pattern. This pattern is intense at first, rapidly receding, but returning to baseline only after several months or years. Thus, ecological research would benefit from a model of disturbance that is inclusive of these events, and in which temporal dynamics could be varied quantitatively in simulation or experimentation (see Ryo et al., 2019). While I did not succeed in finding such a model in the literature, my work in bioacoustics led me to one from sound and music production: the amplitude envelope (Beeman 1998).

## New Research Idea

I describe a model for generating and discussing the temporal dimensions of ecological disturbance based on a popular component of electronic musical instruments – the envelope generator (Deutsch & Deutsch, 1979). This component determines the change over time in its output (typically used to control amplitude), and generally uses four parameters to describe the shape of that change. These parameters, *Attack*, *Decay*, *Sustain*, and *Release*, are described in detail in Figure 1a-b. I will refer to them here as the “ADSR model”. In music, this process is usually triggered by the press of a piano key. However, in its ecological application, I define the ADSR model to be a function of constant parameters describing the intensity of disturbance over time given a single input, the duration of a disturbance event. When the event beings, the function increases in intensity through the duration of the *Attack* stage to its maximum, and decreases through the duration of the *Decay* stage. The intensity then is held at the level of *Sustain* until the event is concluded, at which point the *Release* stage is triggered and continues for its specified duration. Three discrete states are implicitly assumed in the ADSR model: the original state, the maximum amplitude of the signal, and the sustained amplitude of the signal. However, the sustain state can be omitted by setting its value to 0 or 1, and return to the original state can be omitted by continuing the event indefinitely. While there are assumed to be four temporal stages in the model, it can be truncated to only include *Attack* and *Decay* by setting the *Sustain* value to 0, and either the *Attack, Decay, or Release* stage can be omitted by setting their value to 0.

**Figure 1:**
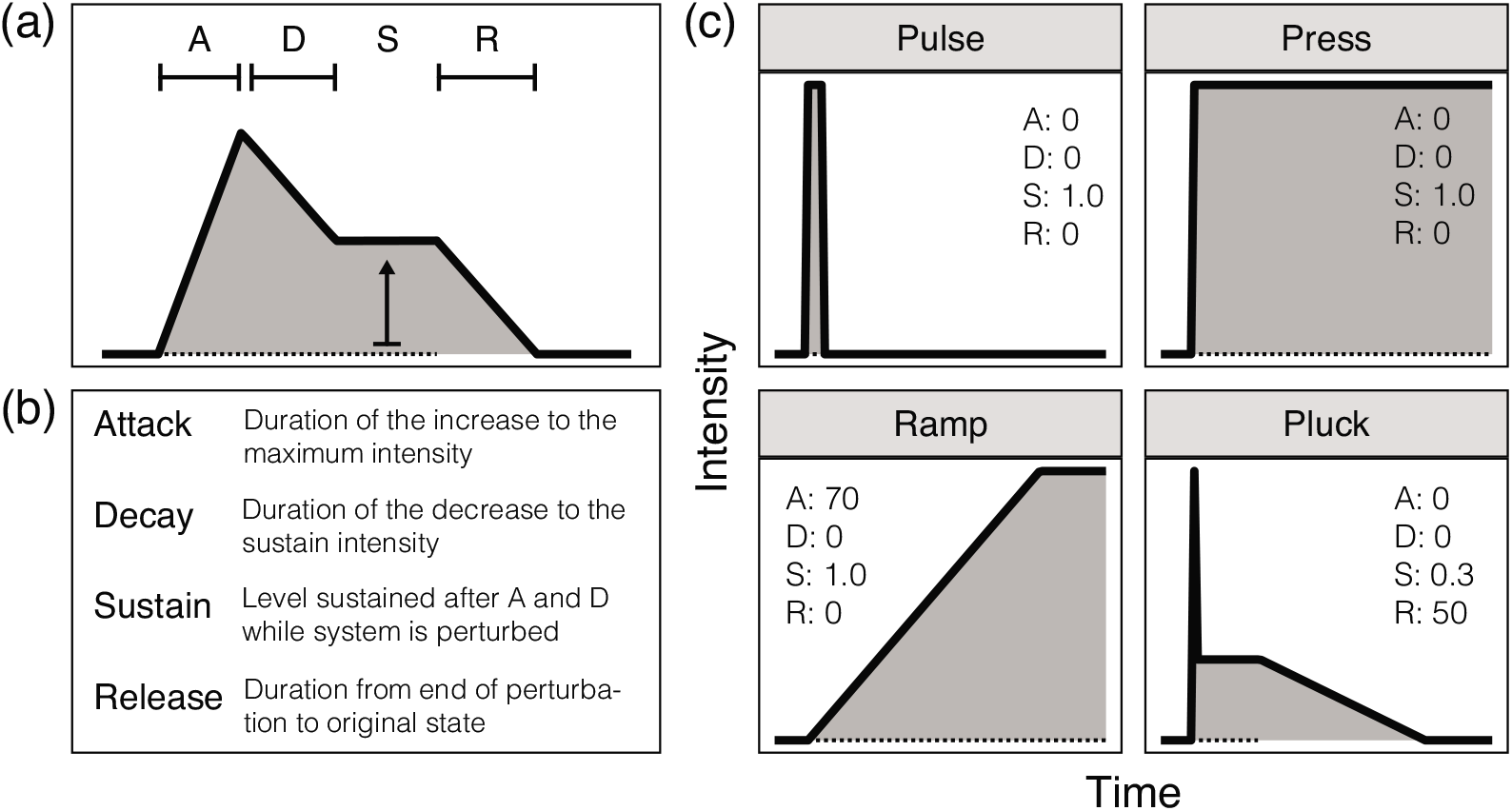
Explanation of the ADSR envelope model (a) with parameter definitions (b); perturbation types described by Lake (2000), and one described in this paper, are reproduced using the ADSR model (c), with parameter values included. Dotted lines indicate the duration of perturbation.

When applied to a simulation of ecological disturbance rather than sound amplitude, the ADSR model is able to describe and generate the features associated with each of type of disturbance described by Lake (2000; Figure 1c). Using the model, one can reproduce “Pulse” disturbances using a short attack and release, and reproduce “Ramp” and “Press” disturbances using continued disturbance with varying attack. In addition to generating common categories of disturbance, the ADSR model should allow researchers to isolate and vary each parameter individually. Based on the dynamics described in many natural systems above, wherein an intense disturbance is followed by a slow return to the original state, I propose a new type of dynamic similar to the “Pluck” of a stringed instrument. This dynamic is intended to imitate the pattern of natural events described above: a brief and intense disturbance, followed by a prolonged recovery to the original state.

The terminology of the ADSR model may only be familiar to some musicians. However, unlike terms with more intuitive but potentially misleading names (e.g., stability and resilience; Donohue et al., 2016), the parameters of the ADSR model have meanings that are strictly quantitative, and so they should hopefully be clearer and more specific for use in discussion among ecologists.

## How to Solve the Question Through the New Idea

Using the R programming environment, I built a simple script for generating ADSR models for use in ecological simulation and modeling (script and demonstration are included in the online supplement). In this implementation, I have assumed that change in intensity is linear for the sake of simplicity. Researchers may wish to use exponential curves for their *Attack, Decay* or *Release* stages; this is entirely feasible, and indeed many synthesizers include this feature. Analog envelope generators are constructed of simple circuits, the length of each stage being determined by varying the rate at which a capacitor should fill. There are numerous ways to emulate this behavior in R or other programming languages, only one of which is described here in the supplement.

To test the effects of various perturbation dynamics on populations, I adapted Jacquet et al.’s (2020) implementation of a logistic growth model with perturbation. This model assumes an intrinsic growth rate that declines as the population approaches a carrying capacity. During each time interval, the population grows – and in Jacquet et al.’s (2020) case a proportion of the population can also be removed. While Jacquet et al. (2020) use this approach to understand the effects of repeated perturbations, I re-configured their model to follow a single perturbation whose intensity (fraction of the population removed) varies over time. To evaluate the utility of the ADSR model, I estimated temporal dynamics of populations responding to the disturbance dynamics described in Figure 1. These show different outcomes depending on the parameters specified: “Press” and “Ramp” disturbances led to near-immediate and eventual extinction, respectively, whereas “Pulse” and “Pluck” disturbances led to immediate and eventual recoveries, respectively (Figure 2).

**Figure 2:**
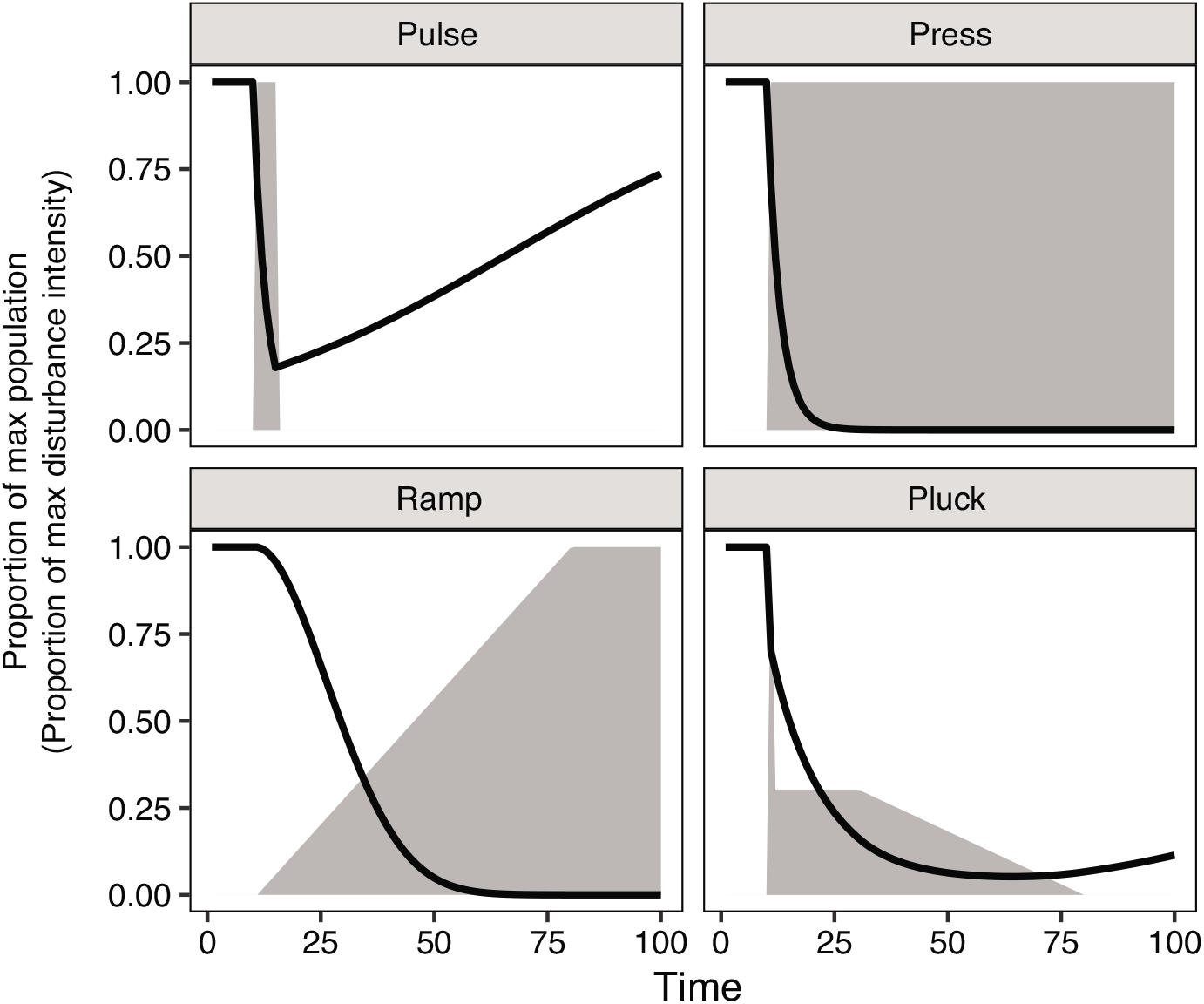
Population effects of disturbance dynamics with types described by Lake (2000), and one described in this paper. The black line represents the proportion of the initial population remaining, while the grey polygon represents the proportion of the maximum disturbance intensity. Disturbance is simulated here by removing 30% of the population per time increment at maximum intensity.

Furthermore, I used the ADSR model to estimate the dynamics of populations with similar growth parameters and overall disturbance, but varying attack and release parameters. I found that increasing attack reduced the impact on the population, and increasing release lessened but lagged the impact (Figure 3a).

**Figure 3:**
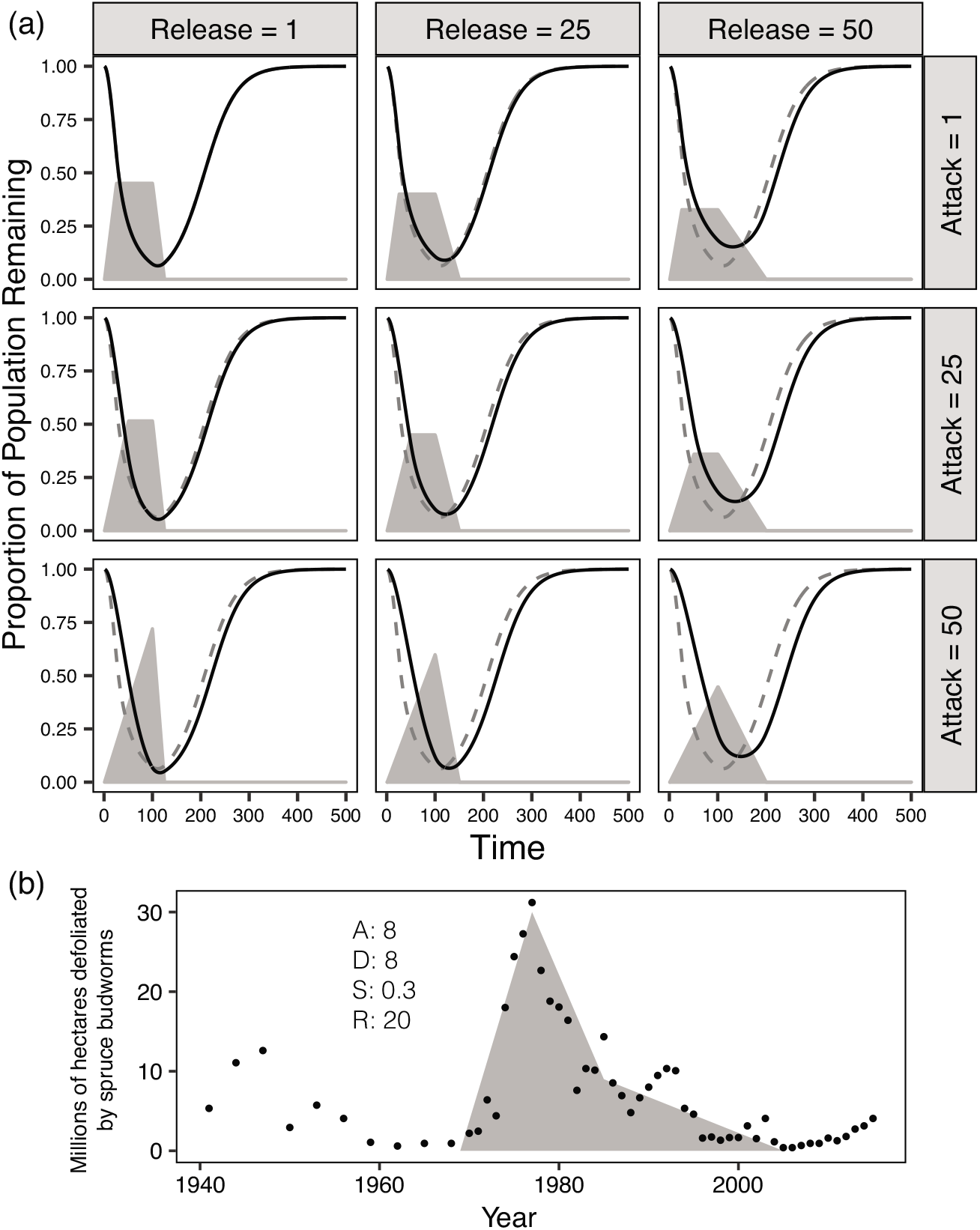
Population effects (a) of disturbance dynamics with varying attack and release parameters (polygons shaded light grey) and their effects on population dynamics assuming a logistic growth model (black line; Jacquet et al. 2020). Dynamics from the upper left panel are included in each subsequent panel as reference (dotted line). Disturbance amplitude values are adjusted to ensure that the area under the curve is approximately equal (models without this adjustment also showed effects of varying attack and release parameters). An example (b) of an ADSR model fit to empirical ecological data, in this case the intensity of a spruce budworm outbreak in North America (re-plotted with permission from Walker & Taylor, 2020). The grey polygon outlines the model with its parameters given at left, and the points represent the sum of hectares showing moderate or greater defoliation for that year.

Lastly, I give a conceptual example of fitting an ADSR model to empirical ecological data (Figure 3b), in this case the impact of spruce budworms on Canadian forests (Walker & Taylor, 2020). Using the ADSR model, I describe this disturbance as having an attack and decay stage of eight years each, with a release stage of twenty years – a dynamic that is a poor fit to any previously described disturbance dynamic. Improvements in fit to these data may be achieved by utilizing a long exponential decay stage in place of the release stage. While the model in the figure is fit arbitrarily, it should be feasible to evaluate and optimize the model’s fit analytically (e.g., Caetano et al., 2010) or through a Bayesian-stochastic approach.

One of the unique implications of “ramp” disturbance with a long *Attack* phase is the potential for species to adapt to their newly altered environment, potentially buffering to some extent the impacts of global climate change (Visser, 2008). To test the effects of various disturbance dynamics on adaptation, researchers may employ evolutionary models such as the multi-locus model developed by Jain and Stephan (2017). A preliminary test of this model, as implemented by Johnsson (2021), showed apparent non-linear behavior in response to varying values of release (see online supplement).

The results presented here ignore many important and relevant factors in ecological and evolutionary dynamics, and are meant primarily to demonstrate the utility of the ADSR model as a tool for future studies. Furthermore, researchers should exercise care in specifying appropriate measurements of disturbance, and especially in separating disturbance from its ecological response. My results are a hint of evidence that varying the dynamics of disturbance, either qualitatively or quantitatively, changes its impact on populations. Experimental studies, and especially empirical studies of long-term ecological data, are needed that examine the resilience of populations and communities to disturbances of varying temporal and also spatial characteristics (Ross et al., 2021).

## Motivation

Much of the impetus for this study comes from personal observations of ecological responses to typhoons on Okinawa Island, Japan. Intense typhoons are brief even on an ecological time scale, but they leave lasting effects that do not return to their original state for years (White et al., 2017). Many animals may perish during the typhoon itself, but more still will find the habitat of their home range no longer suitable, the forest canopy salted over and blown out (Elliott & Nino, 1960) or the coral reef damaged (Nanami & Nishihira, 2002; White et al., 2013). This motivated me to find a model of disturbance that included an intense event with a slow subsequent recovery – a “Pluck” dynamic.

## Acknowledgements

This paper is the result of conversations and thoughts had at the symposium on Ecological Stability at the 66^th^ Annual ESJ meeting in Kobe, and especially with its organizers Sam RP-J Ross and Yuka Suzuki. Ideas in this paper were improved by comments from Claire Jacquet, Martin Johnsson, and Sam RP-J Ross, as well as conversations with members of the Economo Unit, two anonymous reviewers and the editor. The author is supported by subsidy funding to OIST.

## Conflict of Interest

The author declares no conflict of interest.

## Notes

### Competing Interest Statement

The authors have declared no competing interest.

